# Low Field Magnetic Resonance Imaging to Detect Acute Kidney Injury

**DOI:** 10.1101/2025.01.22.634393

**Authors:** Shun Kishimoto, Kazumasa Horie, Nallathamby Devasahayam, Kota Yamashita, Chandramouli Gadisetti, Kazutoshi Yamamoto, Jeffrey R. Brender, James B. Mitchell, Murali C. Krishna, W. Marston Linehan, Daniel R. Crooks

**Affiliations:** Urologic Oncology Branch, Center for Cancer Research, National Cancer Institute, Bethesda, MD, USA; Clinical Cancer Metabolism Facility, Center for Cancer Research, National Cancer Institute, Bethesda, MD, USA; Radiation Biology Branch, Center for Cancer Research, National Cancer Institute; Genepria, Rockville, MD, USA

**Keywords:** Oximetry, Renal Oxygenation, EPR, Acute Kidney Injury

## Abstract

Renal oxygenation is essential for maintaining kidney function. Disruptions in oxygen delivery can lead to renal hypoxia, which can exacerbate kidney injury through multiple pathways, including inflammation, oxidative stress, and ischemia-reperfusion injury. Despite the recognized importance of oxygenation in renal pathology, non-invasive and reliable methods for assessing kidney oxygen levels are limited. Current techniques either lack sensitivity or involve invasive procedures, restricting their use in routine monitoring. Therefore, there is a pressing need for innovative approaches to assess renal oxygenation, particularly in kidney injury. This study evaluated Electron Paramagnetic Resonance (EPR)-based oxygen imaging using the paramagnetic tracer Ox071 to assess kidney oxygen levels in mice with cyclophosphamide-induced kidney injury. Urine pO2 was also assessed as a potential surrogate marker. EPR oximetry accurately measured kidney oxygen distribution, revealing a temporary increase in pO2 post-injury. Urine oximetry, however, did not reliably reflect changes in kidney oxygenation. Furthermore, EPR oximetry provided high-resolution spatial mapping of oxygen levels within the kidney, allowing for a detailed understanding of the impact of hypoxia on renal tissue. EPR oximetry is a promising, non-invasive tool for monitoring renal oxygenation, offering high-resolution mapping and longitudinal assessment. Its ability to provide detailed information about oxygen distribution within the kidney makes it a valuable tool for studying the pathophysiology of renal diseases and for developing novel therapeutic strategies.

Translational Statement: Quantitative spatially resolved measurement of renal oxygenation has the potential to guide clinical decision making in renal disorders such as acute kidney injury. In this study we demonstrate the utility of electron paramagnetic resonance imaging to provide non-invasive and quantitative high-resolution mapping of kidney oxygen concentrations.

## Introduction

The kidney is a highly vascular organ that requires a constant and sufficient oxygen supply to carry out the essential functions of blood filtration, reabsorption of water, and concentration of the urine outflow to an osmolality far exceeding that of the plasma. Renal blood flow constitutes a quarter of the cardiac output, and maintenance of tubular osmotic gradients and solute resorption require large amounts of ATP generated by oxidative phosphorylation in the abundant mitochondria present in the cortical and medullary tubules.^1^ Kidney injury caused by ischemia, cytotoxic chemotherapeutic agents, imaging contrast dyes, certain medications (e.g. NSAIDS and some antimicrobials) can have a significant impact on the perfusion of the kidneys, potentially leading to renal hypoxia.^2, 3^ The relationship between renal oxygenation and kidney function is intricate and may vary depending on the specific disease and individual factors. For instance, conditions like renal artery stenosis^4, 5^, atherosclerosis, or acute kidney injury (AKI)^6^ can lead to decreased blood flow, ultimately reducing the oxygen available to renal cells and contributing to hypoxia.^7, 8^

In the case of AKI, inflammation plays a major role in altering kidney oxygenation. Following injury, the kidneys may be subjected to inflammatory processes, leading to vasoconstriction and decreased blood flow to the injured area, which further reduces oxygen supply to the affected renal tissue.^9^ Persistent inflammation and tubular swelling can sustain hypoxia in the kidneys.^9^ Injured kidney tissue may also experience changes in oxygen utilization. Some cells may revert to glycolysis to sustain energy needs, while others may suffer from mitochondrial dysfunction resulting in decreased oxygen consumption.^10^ These changes can contribute to a relative oxygen deficit in the kidneys, irrespective of sufficient oxygen delivery. Therefore, the overall effect of AKI on local pO_2_ is determined by the balance of reduced oxygen supply and altered oxygen utilization. Kidney oxygenation can remain altered even in later stages of healing from AKI. In response to injury, the kidneys may develop fibrosis and scarring, impairing perfusion and oxygen-carrying capacity and further hindering oxygenation of the affected areas.^11, 12^

To manage kidney injury and prevent further damage, addressing the underlying causes, improving blood flow to the kidneys, and reducing inflammation are crucial.^9, 10^ Early intervention and appropriate treatment are essential to mitigate the effects of kidney injury on oxygenation and overall kidney function. However, currently, there is no established pO_2_ imaging modality available for monitoring renal oxygenation to assess kidney injury and treatment effectiveness.

Electron Paramagnetic Resonance (EPR) imaging with a trityl-based paramagnetic tracer enables non-invasive, quantitative tissue pO_2_ imaging. Studies using the trityl radical tracer, Oxo 63, have successfully mapped hypoxic tumor regions.^13-15^ However, the short signal lifetime of Ox063 can limit access to well-oxygenated areas due to low signal-to-noise ratios. In contrast, Ox071, a deuterated analog of Ox063, demonstrated significantly longer signal lifetimes, providing superior accuracy in estimating pO2 levels in normoxic tissues.^16, 17^ This suggested that Ox071 EPR oximetry could potentially be used to detect acute kidney injury resulting from chemotherapy-related adverse events by monitoring changes in pO_2_ distribution. Unlike other methods, EPR oximetry allows for non-invasive mapping of pO_2_, enabling the detection of focal damage and longitudinal assessment over time.^18^ This study has three primary objectives. First, we investigated our ability to distinguish variations in pO_2_ levels between healthy kidneys and those affected by injury induced through intraperitoneal injections of 200 mg/kg cyclophosphamide. Second, we explored the potential of urine oximetry as a surrogate marker for kidney hypoxia, building on previous findings. Lastly, we aimed to reconstruct high-resolution EPR oximetry images of the kidneys to enhance our understanding of renal oxygenation patterns.

## Methods

### Animals

Female C57BL/6 (B6) mice from the Frederick Cancer Research Center (Frederick, MD) were kept in a controlled environment with regulated circadian rhythms. Experiments were conducted on 8-12 week-old female mice following the Guide for the Care and Use of Laboratory Animals (National Research Council, 1996. All animal experiments were approved by the National Cancer Institute Animal Care and Use Committee (NCI ACUC) and conducted in accordance with NCI ACUC guidelines under the authority of the animal protocol (RBB-159).

The mice were secured in a specialized mount designed for the administration of inhalation gases, including air and anesthesia. For kidney imaging procedures, the animals were positioned within a 25 mm parallel-coil resonator. Anesthesia was administered using 2-2.5% isoflurane, while the mice’s body temperature was carefully maintained at 37±1°C, and their breathing rate was kept between 60-80 breaths per minute. Following a 30-minute stabilization period, a bolus of Ox071, dosed at 1.125 μmoles per gram of body weight, was injected into the tail vein of each mouse.

Kidney oxygenation imaging studies were conducted on the same mouse over three consecutive days, with air (21% O2), carbogen (95% O2 + 5% CO2), and 10% O2+ 90% N2. To study the effect of kidney injury on oxygenation, 200 mg/kg of cyclophosphamide (CPA) was injected intraperitoneally.^19^ Kidney tissue sections were stained with H&E for morphology assessment and with pimonidazole to evaluate oxygenation on day 0 (without CPA), and on days 2 and 30 post-CPA administration. Each group consisted of 8 replicates, and 8 additional mice were used for non-invasive EPRI oximetry at these time points.

### Immunohistochemical analysis

Pimonidazole (60 mg/kg) was administered intravenously 60 minutes before tissue collection. Kidney tissues were frozen, sectioned, and fixed. The sections were blocked and stained overnight at 4°C with Hypoxyprobe™ rabbit antipimonidazole antibody (1: 50; Hypoxyprobe, Inc.) for pimonidazole staining. Fluorescence microscopy and imaging were conducted using a BZ-9000 BIOREVO (KEYENCE). Images were captured with the BZ-9000E viewer at 10□×□magnification and then stitched together to compose a whole image of the sections using the BZ-II Analyzer. Quantification of the pimonidazole-positive fraction was achieved by counting the pixels of the positive area.

### EPR Imaging

EPR-based oxygen imaging relies on the Heisenberg exchange of unpaired spins between O2 molecules and the spin of the trityl radical Ox071, leading to a reduction in the lifetime of the signal.^20^ Due to collisional quenching, the electron relaxation rates R_2_*, R_2,_ and R_1_ of narrow-line trityl radicals like Ox063 and Ox071 increase quantitatively with pO2 levels.^16, 21^ Here we use changes in R_2_* in response to pO_2_ levels calculated from time-domain single point EPR imaging (SPI) in the phase-encoding mode to construct pO_2_ maps of the kidneys.

In SPI, the intrinsic spatial resolution and field of view (FOV) of the image are given by eqs. [1] and [2].

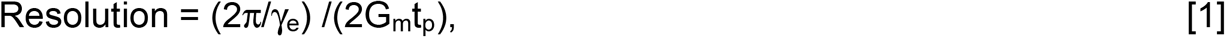

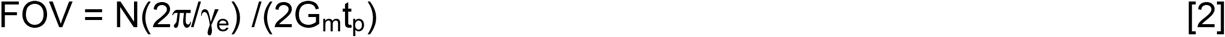

where γ_e_ is electron gyromagnetic ratio, t_p_ is the delay time from the excitation pulse, N is number of phase encoding steps. R_2_* can be calculated from the exponential decay of the signal (S) given by.

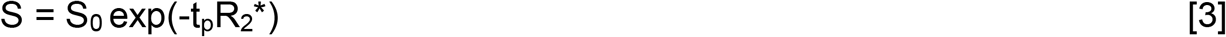

where S_0_ is the intensity at t_p_ = 0. In principle, R_2_* can be collected from a series of EPR image with a range of t_p_ values. In practice, the FOV decreases as t_p_ increases at constant G_m,_ which requires a resampling procedure to align the pixels across the images of different t_p_ values. ^22^

First, one field of view (FOV) is selected as the target. All other images with different tp values are then transformed to match this desired FOV. This transformation process involves two steps: spatial resampling to align the pixels with the desired FOV, and intensity scaling. The intensities of the transformed images are adjusted by multiplying them by a factor Zd, where Z is the ratio of the desired FOV to the original FOV of the image (Z = FOV(desired) / FOV(tp)), and d is the dimensionality of the imaging (d = 3 for 3D imaging). The final R_2_* map is calculate pixel-by-pixel by Eq. 3. This procedure ensures that all images, regardless of their original tp values, are standardized to the same FOV while preserving their relative intensity information, which is crucial for accurate pixel-by-pixel comparison and subsequent calculation of R2* maps across the entire series of images.

In practice, the usable t_p_ range at a single G_m_ is limited because large differences in spatial resolutions lead to edge artifacts in the R_2_* map. These artifacts subsequently appear magnified in the pO_2_ map since pO_2_ is proportional to the change of R_2_*. The t_p_ range can be increased by using different gradient maxima for different t_p_ values, e.g. higher G_m_ for smaller t_p_ values since the resolution depends on the product of t_p_ and G_m_.^22, 23^ We found a three-gradient scan at G_m_ = 1.71, 1.39, and 1.17 G/cm, matrix size =23 x□23□x□23 and a scan time =22 min was adequate to obtain acceptable kidney pO_2_ maps with a negligible change in resolution across the tp range (1.5382 ±0.0008)..

EPR imaging of mice was carried out using a custom-built FT-EPR imaging system operating at 300 MHz.^22^ Time-domain EPR signals were recorded at ambient temperatures through a single excitation pulse sequence with pulse width = 70 ns (flip angle ∼ 50°), repetition time TR = 8 μs, FID signal averages = 4000, number of phase cycles = 4, and number of sampled points = 1000, with a dwell time of 5 ns. The 3D EPR imaging data were acquired in single-point imaging (SPI) mode which is a phase encoding approach, utilizing a 23L□×□23□×□23 Cartesian grid. Raw k-space data underwent correction for any DC shifts, and high-frequency noise was filtered using a Tukey window (r = 0.7). Image reconstruction was conducted through fast Fourier transformation after zero-filling the k-space matrix to 64□x□64 x□64 or higher. Oxygen imaging was performed using a multi-gradients scan at three gradient maxima (Gm) = 1.71, 1.39, and 1.17 G/cm, with a scan time of approximately 22 minutes. The R2* values were computed from the exponential decay of pixel intensity (see Results section). Tissue pO_2_ maps were derived from R2* using previously established linear relationships between R2* and pO_2_. Higher resolution scans were conducted with a matrix size of 25□× □25L× □25 and Gm values of 1.87, 1.52, and 1.28 G/cm, with a scan time of approximately 28 minutes.

### Calculation of Kidney and urine pO_2_

Kidney pO_2_ imaging datasets utilizing a matrix size of 23 × 23 × 23 and gradient strengths (Gm) of 1.71, 1.39, and 1.17 G/cm were reconstructed at tp = 405 nsec, yielded a spatial resolution of 1.32 mm, further enhanced digitally to 0.48 mm. High-resolution kidney probe distribution imaging dataset utilizing a matrix size of 25L×□25□×□25 and Gm values of 1.87, 1.52, and 1.28 G/cm was reconstructed at tp = 750 nsec resulted in a spatial resolution of 0.84 mm, further digitally improved to 0.17 mm. Subsequently, high-resolution kidney pO_2_ imaging was derived from the same dataset, reconstructed at tp = 520 nsec, achieving a spatial resolution of 1.06 mm, further digitally improved to 0.22 mm. Throughout the data processing, 3D rotations were applied to align the kidney direction for visualization. To measure urine pO_2_, the ureter was initially identified as a cord-like structure connecting the kidney and bladder based on the Ox071 signal intensity images used in the kidney pO_2_ imaging datasets. Subsequently, the ureter region displaying a homogeneous distribution of pO_2_ values proximal to the kidney was chosen as the urine region. This selection was made under the assumption that there is minimal oxygen exchange between the urine and the ureter wall in that particular region.

### Statistical Analyses

A one-way ANOVA followed by Dunnett’s multiple comparisons test was conducted for the histological assessment (Fig. 3D), while paired t-tests were used to evaluate longitudinal changes in pO□ (Fig. 4D and 5D). All statistical analyses were performed using GraphPad Prism (version 10.0.0) for Windows, GraphPad Software, Boston, Massachusetts, USA (www.graphpad.com). Statistical significance was defined as P < 0.05 and P < 0.01.

## Results

### EPR Oximetry Maps Renal Oxygen Distribution with High Spatial Resolution

The median intensity map (S_0_ averaged across all tp values) of a representative slice of the 3D image is shown in Fig. 1A. A total of 12 tp values were used to construct the R_2_* map with four t_p_ values from each G_m_ as shown in Fig. 1B. The intensity map indicates a high accumulation of Ox071 in the kidneys, which appeared significantly brighter than surrounding tissues (Fig. 1A). The spatial resolution in the pO_2_ map is sufficient to detect the internal architecture of the kidneys, with the kidney cortex clearly delineated from the medulla in Fig. 1C.

**Figure 1.**
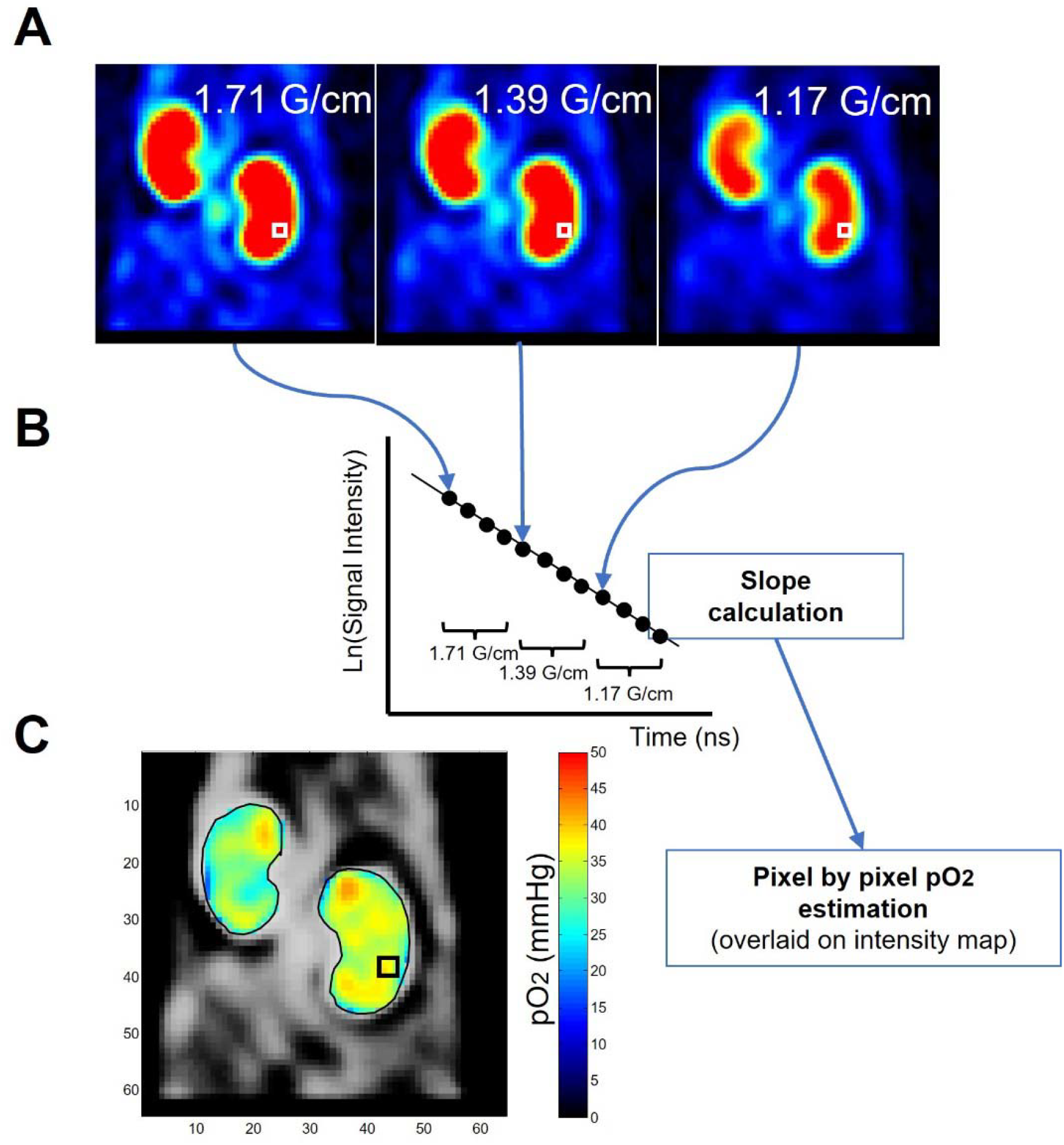
Scheme for estimating pO2 using EPR oximetry involves comparing 12 signal intensity images acquired with three different gradients (four delay time point images per gradient) **(A)** on a pixel-by-pixel basis. The exponential decay of each pixel’s signal intensity (R2* relaxation) **(B)** allows for a direct interpretation of the pO2 value in that pixel **(C)**.

To enhance image resolution for kidney pO2 mapping, we increased the number of gradient steps and utilized FID data from later delay times. We employed 25 gradient steps for all axes and a 10 nsec delay time for probe intensity mapping, resulting in a reduced FOV of 20.9 mm and improved spatial resolution of 0.84 mm (Figure 2A). The heterogeneous signal distribution in the image reflects the kidney’s structural complexity, with high-intensity areas corresponding to the renal cortex and low-intensity regions indicating the medulla and pelvis. Using the same dataset, we reconstructed the pO2 map using twelve delay time points (520-850 nsec) and three maximum gradients (1.87, 1.52, and 1.28 G/cm). The resulting pO2 map (Figure 2B) has a FOV of 26.4 mm and 1.06 mm spatial resolution, showing higher pO2 values in the peripheral cortex and lower values in the core medulla, consistent with established pO2 gradients from previous studies.

**Figure 2.**
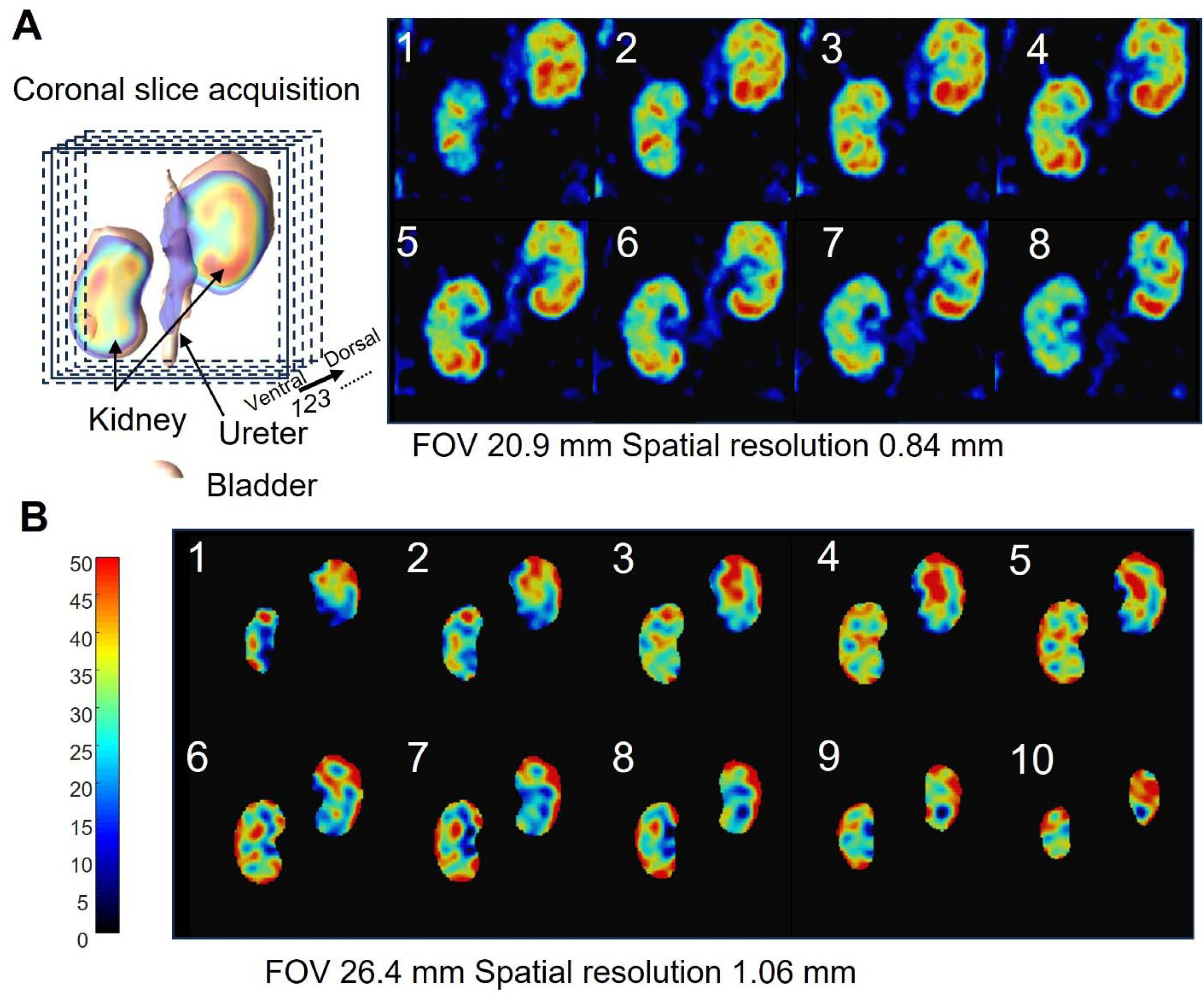
High-resolution imaging of kidney EPR images. **(A)** Slice direction for EPR images is illustrated in the left panel, while the right panel shows high-resolution EPR signal intensity images with a spectral resolution of 0.84 mm. **(B)** High-resolution EPR pO2 images with a spectral resolution of 1.06 mm.

**Figure 3.**
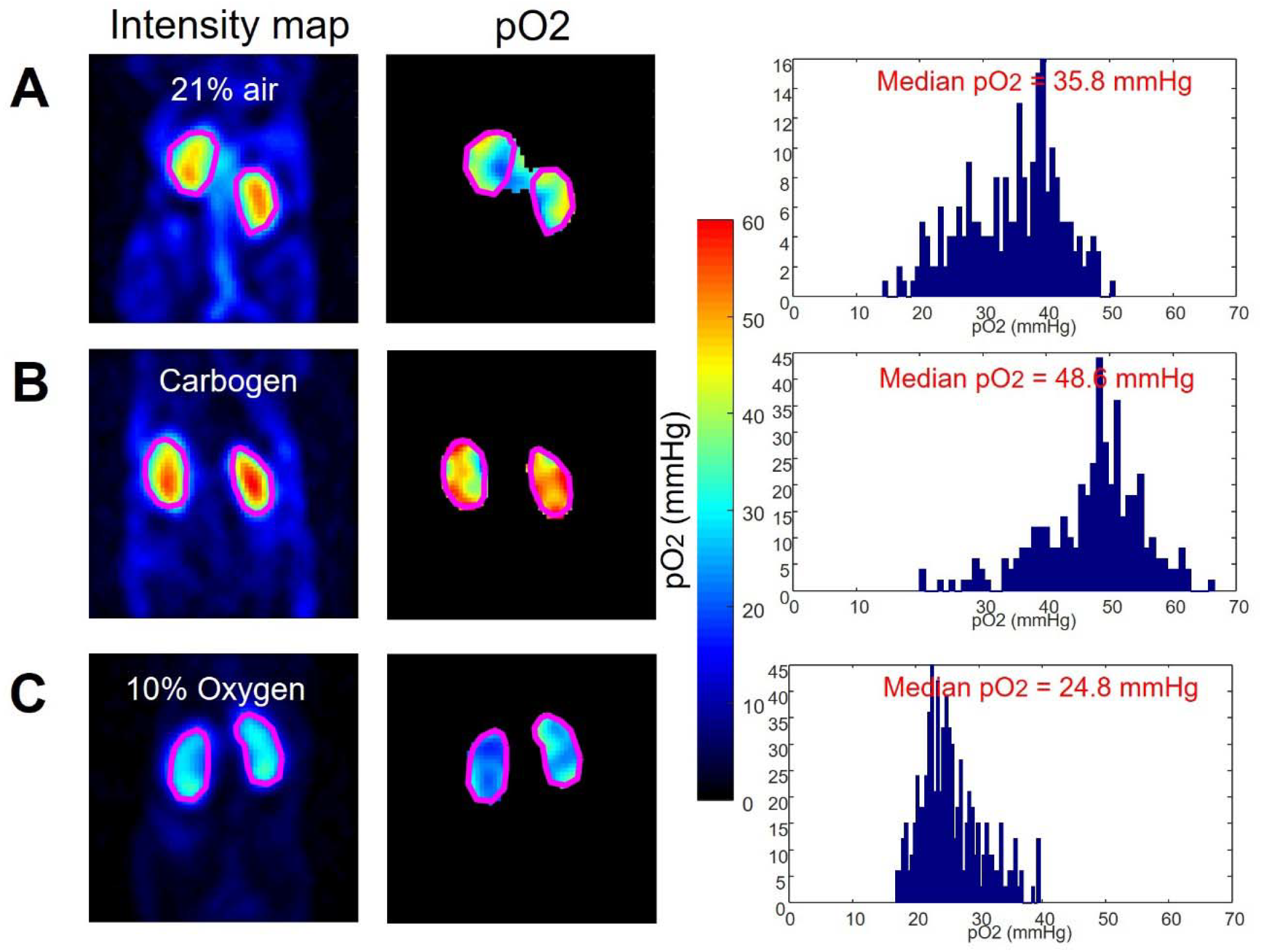
EPR pO_2_ imaging of renal oxygenation in mice exposed to air **(A)**, carbogen **(B)**, and 10% oxygen **(C)**. In each case **(A-C)**, the left, center, and right panels display the EPR intensity map, EPR pO_2_ map, and histogram of intra-kidney pixels, respectively. To mitigate the residual effects of inhalation gas, the experiment was conducted at one-day intervals over three consecutive days.

**Figure 4.**
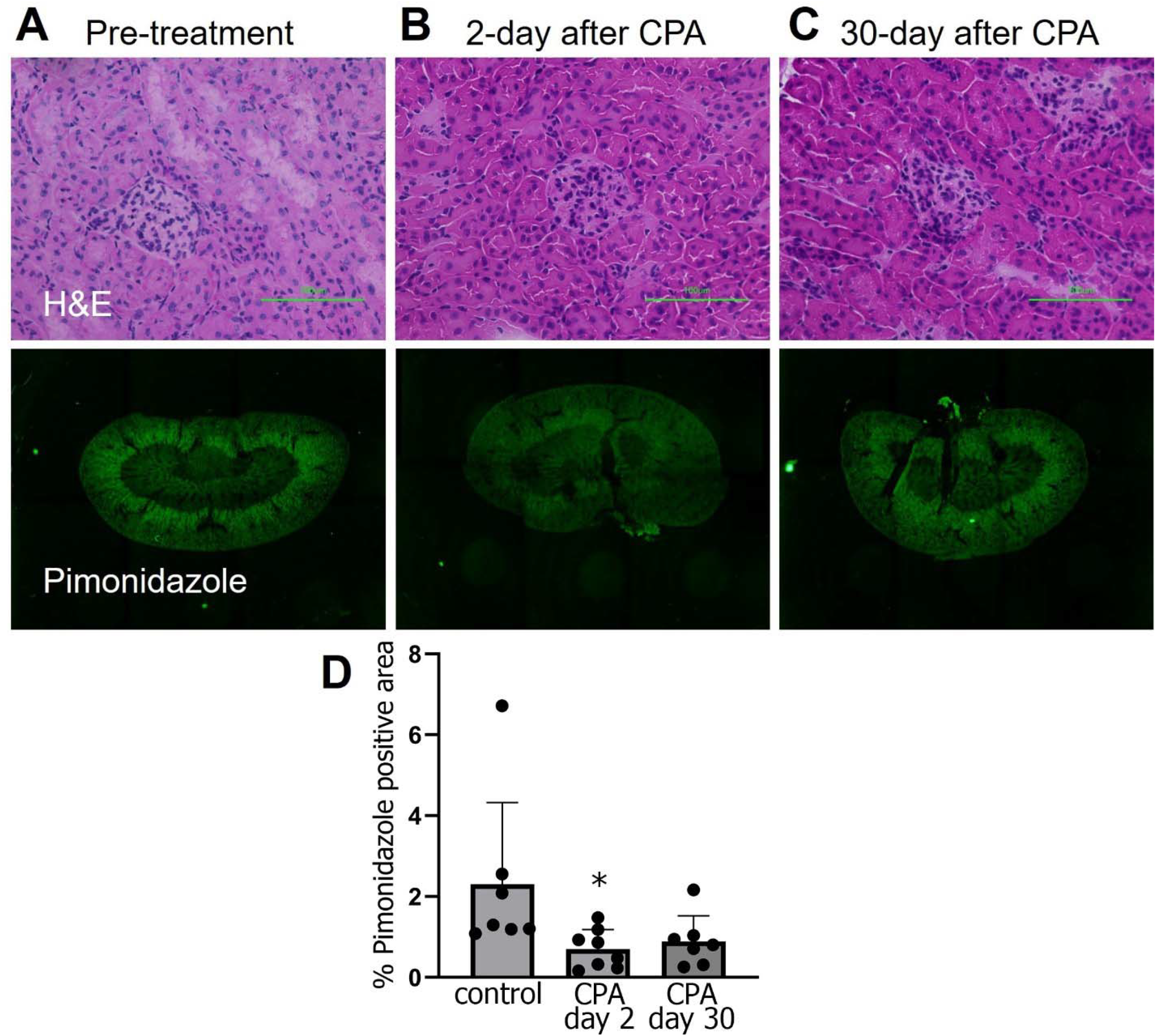
Cyclophosphamide (200 mg/kg) induced alterations in the kidney assessed using H&E and pimonidazole staining. **(A)** In the control group, H&E staining revealed a normal tissue structure, and pimonidazole staining showed a distinct hypoxia gradient between the medullary and cortex regions. **(B)** On day 2, H&E staining indicated tubular swelling and loss of brush border, and pimonidazole staining revealed a global reduction in hypoxic regions. **(C)** By day 30, although morphological changes resembled those observed on day 2 in H&E stained tissue, the hypoxic regions were observed to have recovered in pimonidazole stained tissue. **(D)** Comparison of the percent pimonidazole stained area performed between samples (Control group: n = 7, Day 2 after treatment group: n = 8, Day 30 after treatment group: n = 7).

### EPR Oximetry Detects Dynamic Kidney Oxygenation Changes Under Different Respiratory Conditions

Validation of EPR oximetry’s ability to quantify changes in kidney tissue oxygenation was achieved through controlled studies involving mice exposed to a range of oxygen levels. Mice were exposed to a series of respiratory conditions, including normoxia (21% oxygen in air), hyperoxia with carbogen (95% oxygen, 5% carbon dioxide), and hypoxia (10% oxygen, 90% nitrogen). Carbogen, with its 5% carbon dioxide content, was specifically chosen to induce vasodilation^24^, counteracting the potential vasoconstriction resulting from high-concentration oxygen inhalation^25^, which could otherwise decrease oxygen delivery.^26^

The impact of inspired oxygen concentration on kidney oxygenation is illustrated in Figure 3, where panels A, B, and C show intensity maps, pO2 distributions, and histograms for normoxic (21% air), hyperoxic (carbogen), and hypoxic (10% oxygen) conditions. Consistent with expectations, carbogen inhalation elevated renal pO_2_ (median 48.6 mmHg) relative to air (35.8 mmHg), while 10% oxygen reduced it (24.8 mmHg). This systematic variation in kidney oxygenation levels demonstrates the technique’s capacity to accurately reflect changes in tissue oxygen concentration attributed to differences in inhaled oxygen levels.

### Longitudinal EPR Study Captures Kidney Oxygenation Shifts After CPA Treatment

We next explored the potential of EPR oximetry to measure alterations in kidney tissue oxygenation in response to a chemotherapy-induced model of kidney injury. In prior studies we observed a transient reduction in tumor hypoxia in pancreatic cancer models treated with evofosfamide, a hypoxia-activated prodrug that generates nitrogen mustard bromo-iso-phosphoramide selectively in hypoxic cells.^27, 28^ The study linked reoxygenation after treatment not to changes in perfusion but to reduced oxygen consumption, attributed to either cell death or cell cycle arrest. Following a similar rationale, we opted to use high dose cyclophosphamide (CPA), the most widely employed nitrogen mustard-type alkylating agent, to induce kidney injury and examine potential changes in renal oxygenation. Given its modest effects on kidney function, we administered CPA at a single high dose of 200 mg/kg through intraperitoneal injection, a dose that has been previously shown to induce histological alterations in mouse kidneys.^19^

Figure 4 shows the histological assessment of mouse kidneys following CPA treatment. Images of H&E and pimonidazole staining are provided for untreated mice (Figure 4A), and for mice 2 days (Figure 4B) and 30 days (Figure 4C) after CPA treatment. Significant tubular swelling caused by CPA was histologically visible in Figures 4B and 4C. Pimonidazole staining was notably more intense in cortical regions relative to medullary regions, consistent with the higher cellularity and perfusion of the renal cortex compared to the medulla. Similar to previous studies on the effects of nitrogen mustard agents on tumor hypoxia^27^, the pimonidazole staining of the kidney hypoxic area fraction decreased temporarily on Day 2 (*P*=0.038, pre-treatment vs. CPA day 2), and by day 30 was not statistically different from the controls (P=0.078), despite the sustained histological changes in renal tubular architecture.

To validate the ability of EPR oximetry to non-invasively detect changes in kidney pO_2_, eight mice were treated with 200 mg/kg CPA and underwent repeated EPR oximetry scans. Figures 5A-C present representative EPR oximetry results before treatment, 2 days after treatment, and 30 days after treatment, respectively. The left column shows the probe signal intensity map defining the kidney region, the center column shows the corresponding pO_2_ map, and the right column illustrates the histogram of pO_2_ values for pixels in the region of interest (ROI). Figure 5D depicts changes in pO_2_ for each subject as dot plots, with the average values of all data summarized in bar plots. The results are in excellent agreement with the histological assessment, showing a reduction in hypoxia at 2 days after treatment and a return to baseline oxygen levels at day 30.

**Figure 5.**
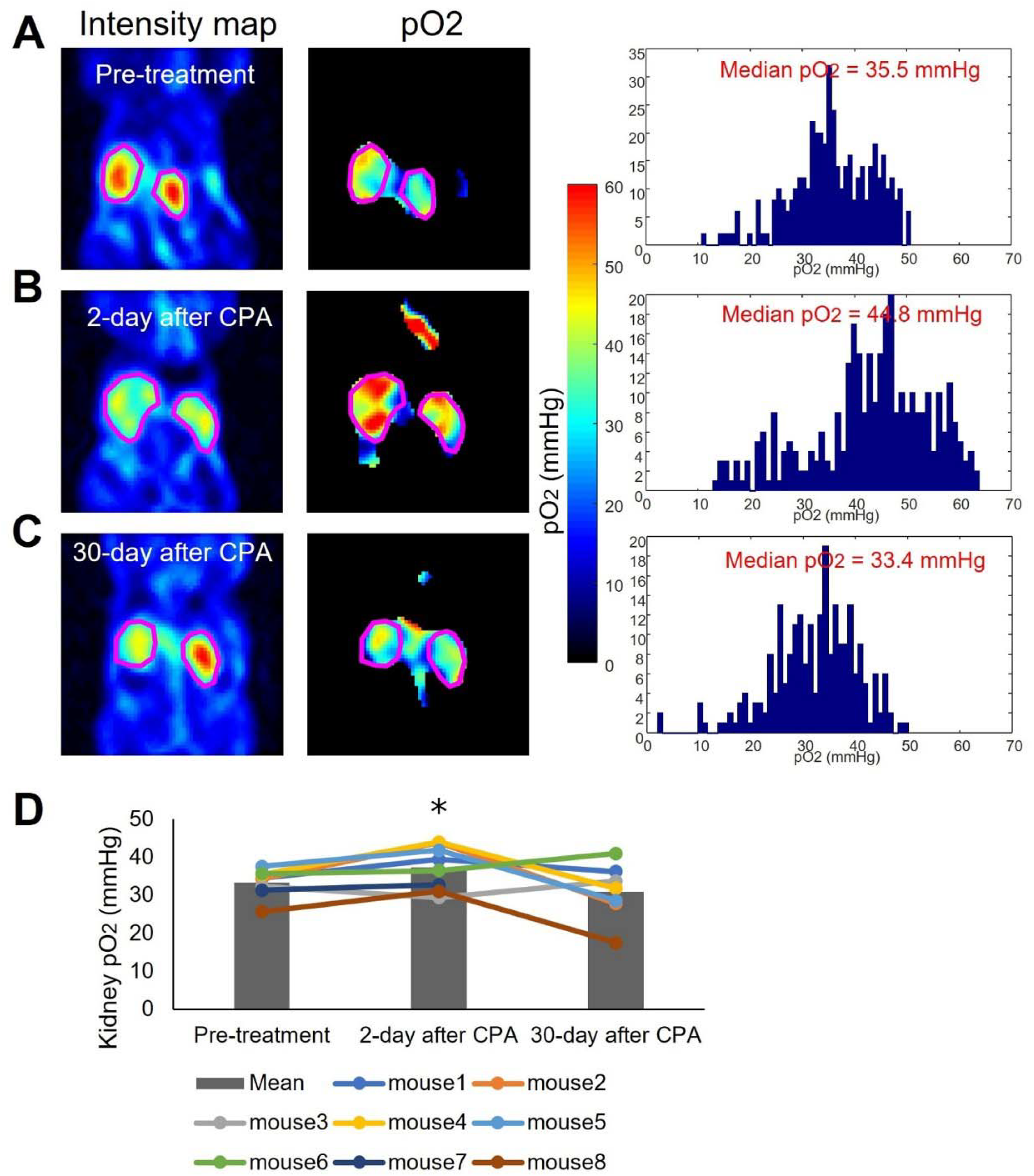
Alterations in kidney pO2 following cyclophosphamide treatment (200 mg/kg). **(A)** represents control kidneys, **(B)** represents kidneys observed 2 days after treatment, and **(C)** represents kidneys observed 30 days after treatment. In each case **(A-C)**, the left, center, and right panels display the EPR intensity map, EPR pO2 map, and histogram of intra-kidney pixels, respectively. **(D)** illustrates the changes in median pO2 within the kidney.

### Urinary pO2 Fails to Reliably Reflect Kidney Tissue Oxygenation After Injury

We next compared urine pO_2_, a surrogate marker for kidney hypoxia under investigational use, with direct kidney tissue oxygenation measurements obtained through EPR oximetry.^29, 30^ The use of urinary pO_2_ as a biomarker for kidney oxygenation has been hindered by two key questions: first, whether sensors can accurately measure urinary pO_2_ under clinical conditions, and second, whether urinary pO_2_ truly reflects medullary tissue oxygenation. The quantitative nature of EPR oximetry effectively resolves the first concern by providing precise, absolute pO_2_ measurements. This allows us to focus on the critical second question: does urinary pO2 accurately represent medullary tissue pO_2_? To address this, we leveraged EPR oximetry’s spatial resolution to simultaneously measure and compare renal tissue pO2 with urine pO2 in the ureter. This approach enables a direct evaluation of the relationship between kidney tissue oxygenation and urinary pO2, potentially validating or challenging the use of urine pO2 as a non-invasive indicator of kidney oxygenation.

Changes in urine pO_2_ after treatment were plotted in Figure 6D, mirroring the plot in Figure 5D for kidney pO_2_. In the data from mice before treatment, kidney pO_2_ and urine pO_2_ displayed an excellent linear correlation. However, in mice data from 2 days post-treatment, the correlation weakened slightly, and it nearly disappeared in mice data from 30 days post-treatment. Notably, urine oximetry did not exhibit the temporary reduction of hypoxia observed in kidney histological assessment and kidney oximetry. This suggests that urine pO_2_ may not be an accurate surrogate biomarker for kidney pO_2_ when the kidney tubules are damaged by cyclophosphamide.

**Figure 6.**
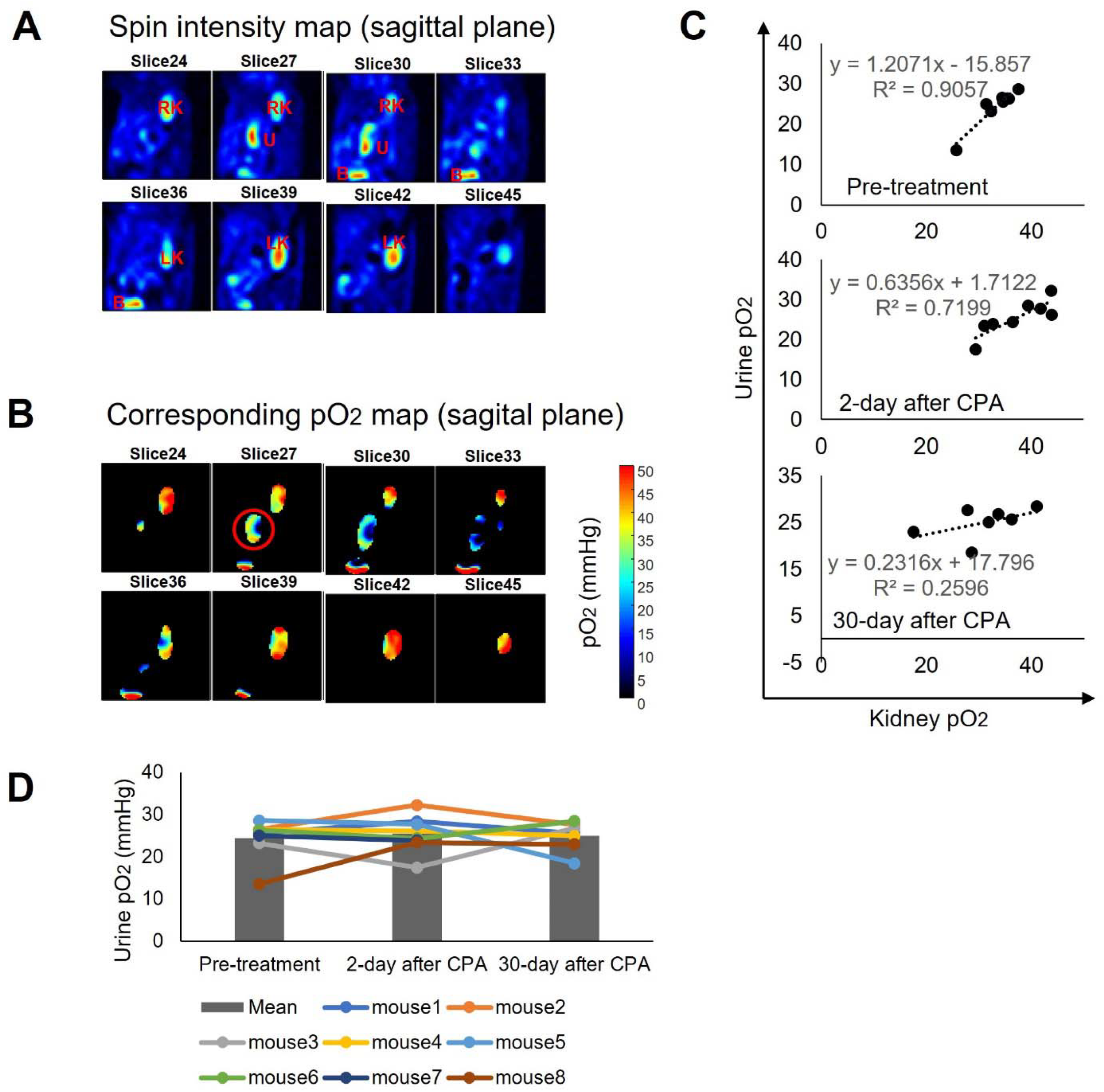
Changes in urinary pO2 after cyclophosphamide treatment (200 mg/kg). **(A)** Signal intensity maps are presented to identify the ureter filled with Ox071-containing urine, with RK, LK, U, B denoting the right kidney, left kidney, ureter, and bladder, respectively. **(B)** Corresponding pO2 maps, clearly indicating urine in the ureter with homogeneous pO2 values. **(C)** Correlation between kidney pO2 and urine pO2 in each group. **(D)** Changes in median pO2 within the kidney.

## Discussion

Given that many chemotherapeutics have the potential to harm the kidneys and kidney hypoxia induced by chronic renal diseases can further contribute to renal damage, there is a critical need to monitor kidney oxygenation changes quantitatively and longitudinally through less invasive imaging methods. Currently, quantitative *in vivo* pO_2_ measurement depends on needle electrode measurements, which, though effective, are too invasive for human studies.^15, 31^ Alternative approaches such as Near-Infrared Spectroscopy (NIRS) and Blood Oxygen Level-Dependent MRI (BOLD-MRI) have been developed and investigated for their efficacy in assessing renal oxygenation.^32^ NIRS is limited by the limited penetration depth of NIR light, and BOLD-MRI lacks quantification.^33-35^ More recently, Dynamic Nuclear Polarization (DNP)-MRI has shown utility in measuring spatial differences in renal oxygenation in diabetic mice, though the methodology is technically challenging and relies on external standards for quantification.^36^ Another alternative avenue explored has been the use of urine pO_2_ as a surrogate marker for kidney pO_2_.^29, 30^ Despite such exploration, the reliability of this method has not been conclusively confirmed. Therefore, the primary objective of this study was to validate the effectiveness of EPR imaging for non-invasive renal oximetry.

EPR oximetry is considered the “Gold Standard” for quantitative tissue oximetry measurements.^37^ We measured the average healthy kidney pO2 as 37.8 ± 3.6 mmHg, consistent with previous studies reporting 20-30 mmHg in the medulla and 50 mmHg in the cortex of mice. ^38, 39^ This concordance with previous studies validates the reliability of EPR for renal oximetry. Our study demonstrates that longitudinal EPR pO2 imaging successfully detected changes in kidney oxygenation following cyclophosphamide treatment, including a temporary increase in pO2 levels. These findings correlated well with histological evaluations using pimonidazole staining.

Cyclophosphamide at doses lower than that used in the present study have been utilized widely as an antiproliferative agent in the treatment of cancer^40^, to control inflammatory diseases including those of the kidney^41^, and in the prevention of graft-versus-host disease during bone marrow and organ transplantation.^42^ Though not considered to be a nephrotoxic agent when administered at therapeutic doses, cyclophoshamide therapy is known to induce a water excretion deficit (syndrome of inappropriate antidiuretic hormone secretion, SIADH) characterized by urine hyperosmolality, decreased plasma osmolality and hyponatremia ^43^. The delayed onset of this antidiuretic effect following cyclophosphamide administration is consistent with the passage of alkylating metabolites of cyclophosphamide through the renal tubules^44^, suggesting a direct renal tubular mechanism of action. Our observation of clear and lasting histopathological effects on renal tubular structures are suggestive of damage to multiple nephron components both two and 30 days following high dose cyclophosphamide administration. This could be due either to direct nephrotoxicity or to systemic toxicity such as cellular lysis in distant organs resulting in glomerular and/or tubular damage analogous to that which occurs during tumor lysis syndrome and crush injury.

Our study evaluating urine pO2 as a potential surrogate marker for kidney oxygenation revealed important limitations of this approach. The linear correlation between urine and kidney pO2 weakened significantly 30 days post-treatment, suggesting urine pO2 may not accurately reflect kidney oxygenation in kidneys damaged by high dose cyclophosphamide. This discrepancy likely results from reduced oxygen exchange between urine and medullary tissue due to tubular damage, as confirmed by histological assessments. These results caution against relying on urine pO2 as a proxy for kidney oxygenation in cases of renal injury. However, the extent to which the ureteral epithelium was damaged following high dose cyclophosphamide administration was not evaluated in our study. Hemorrhagic cystitis due to accumulation of the cyclophosphamide-derived metabolite acrolein is a well known toxicity of this agent ^46, 47^, and pyelitis and ureteritis following cyclophosphamide treatment have been reported ^47, 48^. Thus, reduced integrity of the upper urothelial tracts due to cyclophosphamide toxicity may have contributed to the loss of correlation between renal and ureteral pO2 following treatment.

Our investigation into high-resolution EPR imaging demonstrated its ability to visualize the oxygen gradient between kidney cortex and medulla, as well as the heterogeneous probe distribution in healthy kidneys. This pO2 gradient stems from the uneven distribution of renal blood flow ^49^. The cortex receives the majority of blood flow, while only 10-15% is directed to the medulla to maintain osmotic gradients and enhance urinary concentration. As a result, the medulla typically has lower oxygen tension than the cortex. In conditions such as diabetes and chronic kidney diseases, studies show that medullary oxygenation levels are further reduced compared to cortical levels, with a more pronounced decrease than in healthy individuals ^50, 51^. This heightened vulnerability of the medulla is due to its limited blood supply and higher oxygen consumption, making it susceptible to hypoxic damage, which can lead to renal interstitial fibrosis and disease progression ^52^. Therefore, the ability to assess the oxygen tension gradient between medulla and cortex is clinically significant. The renal oxygen gradient serves as a potential biomarker for detecting early or more severe decreases in medullary oxygen levels relative to cortical levels, which could indicate kidney dysfunction or disease progression.

## Disclosure

The authors declare no competing interests

## Acknowledgements

The authors would like to thank Dr. Christopher Kanakry for thoughtful comments on this manuscript. This work was supported by the Intramural Research Program of NCI-CCR (ZIC BC011932, 1ZIABC010476-22). The content of this publication does not necessarily reflect the views or policies of the Department of Health and Human Services, nor does mention of trade names, commercial products, or organizations imply endorsement by the U.S. Government.

## Notes

### Competing Interest Statement

The authors have declared no competing interest.

